# Deep Visual Proteomics advances human colon organoid models by revealing a switch to an *in vivo*-like phenotype upon xenotransplantation

**DOI:** 10.1101/2024.05.13.593888

**Authors:** Frederik Post, Annika Hausmann, Sonja Kabatnik, Sophia Steigerwald, Alexandra Brand, Ditte L. Clement, Jonathan Skov, Theresa L. Boye, Toshiro Sato, Casper Steenholdt, Andreas Mund, Ole H. Nielsen, Kim B. Jensen, Matthias Mann

## Abstract

Intestinal epithelial damage predisposes to chronic disorders like inflammatory bowel disease. The organoid model allows cultivation, expansion and analysis of primary intestinal epithelial cells and has been instrumental in studying epithelial behavior in homeostasis and disease. Recent advances in organoid transplantation allow studying human epithelial cell behavior within the intestinal tissue context. However, it remained unclear how organoid transplantation into the colon affects epithelial phenotypes, which is key to assessing the model’s suitability to study human epithelial cells. We employed Deep Visual Proteomics, integrating AI-guided cell classification, laser microdissection, and an improved proteomics pipeline to study the human colon. This created an in-depth cell type-resolved proteomics resource of human intestinal epithelial cells within human tissue, *in vitro* organoids, and the murine colon post-xenotransplantation. Our findings reveal that *in vitro* conditions induce a proliferative organoid phenotype, which was reversible upon transplantation and adjustment of organoid culturing conditions.

## Introduction

The intestinal epithelium forms an integral barrier between the intestinal lumen, filled with microbiota and dietary components, and the lamina propria containing immune cells and fibroblasts. Continuous proliferation of epithelial stem cells located within the epithelial crypts ensures constant replenishment of intestinal epithelial cells (IECs). As stem cell progeny move towards the crypt top, they cease to divide and differentiate terminally, establishing a heterogeneous continuum along the crypt axis. These terminally differentiated IECs include absorptive colonocytes, mucus producing goblet cells, and hormone secreting enteroendocrine cells, which all perform key functions in intestinal physiology^1^.

Epithelial maintenance is key for human health and requires tight molecular regulation balancing cell proliferation, differentiation and death. Murine models have provided substantial mechanistic insights into these intricate relations. There are, however, clear differences between the human and mouse, e.g. unique cell types identified in the human intestine^2^, highlighting the need for human models. Addressing mechanistic questions in humans *in vivo* is challenging, and organoids^3–5^ have emerged as an important model system to culture primary human cells and allow experimental manipulation. Human intestinal organoids have provided insights into e.g. cell fate choices, with applications in molecular medicine, drug testing and cellular therapies^1,6–9^. Conventional organoid culture features epithelial cells, but lacks other cell types present in the intestinal mucosa, such as immune cells and fibroblasts^1^. To address this limitation, orthotopic transplantation models have recently been developed^10,11^. They enable the transplantation of wild-type or genetically engineered mouse or human organoids into the murine colon to mechanistically dissect epithelial phenotypes within the mucosal microenvironment, which was previously only possible in mouse models. Furthermore, autologous transplantation of organoids into patients with impaired IEC phenotypes has great therapeutic potential in regenerative medicine, e.g. for inflammatory bowel disease (IBD) and short bowel syndrome^6,8^. This tractable xenotransplantation system enables the assessment of human IEC phenotypes in the mucosal microenvironment^11,12^, but we still only have limited knowledge on how well human IECs transplanted into the murine colon recapitulate human IECs *in vivo*.

Fully leveraging the potential of human organoids requires in-depth characterization and validation of organoid models^1,13^, which necessitates an accurate reference data set of their *in vivo* counterpart. Such a resource could guide future evaluation of disease-related changes, cellular and disease markers, and improvement of *in vitro* model systems. An accurate assessment of cellular phenotypes should account for their spatial context, especially in delicately organized tissues like the colon mucosa. Spatial transcriptomics and fluorescent *in situ* hybridization (FISH)-based techniques have provided valuable insights into the cellular heterogeneity of the colon^14,15^. These approaches, however, require pre-defined target panels and are biased by current knowledge. Single cell RNA-sequencing (scRNAseq), facilitates in-depth characterization of cellular phenotypes^16–18^, but lacks spatial information. Typically, it also requires cellular dissociation and long enrichment protocols, which in itself can impact epithelial phenotypes^19^. In the context of organoids, scRNAseq has been used to assess cellular composition^5,20^, but in-depth phenotypic benchmarking including direct comparison to the *in vivo* counterparts remains limited, especially for the human colon.

Recent studies suggest that deep and sensitive proteomics provides more robust readouts for cellular states than transcriptomes, while directly pinpointing functional consequences of perturbation-induced changes^21,22^. The sensitivity of proteomics has advanced massively in the last decades from the quantification of a few thousand proteins from milligrams of input material in the beginning of the millennium to comparable numbers from single cells to date^22–24^. However, so far none of these methods have reached substantially complete coverage of cell type-specific proteomes. To address this, we here substantially further develop our Deep Visual Proteomics (DVP)^25^ pipeline, which employs high-resolution fluorescence imaging, AI-guided cell segmentation and classification, single-cell isolation by laser capture microdissection, and high-sensitivity proteomics. To date, the conventional DVP pipeline generally yielded up to 5,000 proteins by combining a few hundred contours of single cell contours of the same type^25^. Our improved workflow using low flow gradients and the novel Orbitrap Astral analyzer^26^, improved proteome coverage substantially, from even fewer contours. This allowed us to build a spatial proteome atlas of the human colon mucosa with unprecedented cell type-specific proteome depth. Importantly, the increased depth of protein quantifications at decreased input amounts enabled us to robustly and accurately benchmark human colon organoids grown *in vitro* and transplanted into the murine colon.

Our findings reveal that despite a robust correlation between *in vitro* and *in vivo* proteomes, IECs grown as organoids *in vitro* display high proliferation and low functional signatures. Strikingly, this is reverted upon xenotransplantation, rendering xenotransplanted human IECs a valuable tool to dissect human IEC phenotypes and illustrating that organoids retain their ability to reform colonic epithelium. Combined with iterative, proteomics guided improvements in organoid cell culture conditions this is a promising approach in regenerative medicine.

## Results

### DVP enables in-depth spatial proteomic profiling of cellular populations in the human colon

The assessment of human organoid models requires the determination of the *status quo* of the human colon mucosa. We made use of DVP (Fig. 1A) to generate a high sensitivity spatial proteome atlas of the human colon mucosa and analyze organoid models. In total, we analyzed 11 human colon tissue sections, 15 sections of organoids *in vitro*, and 50 sections of transplanted organoids.

**Figure 1:**
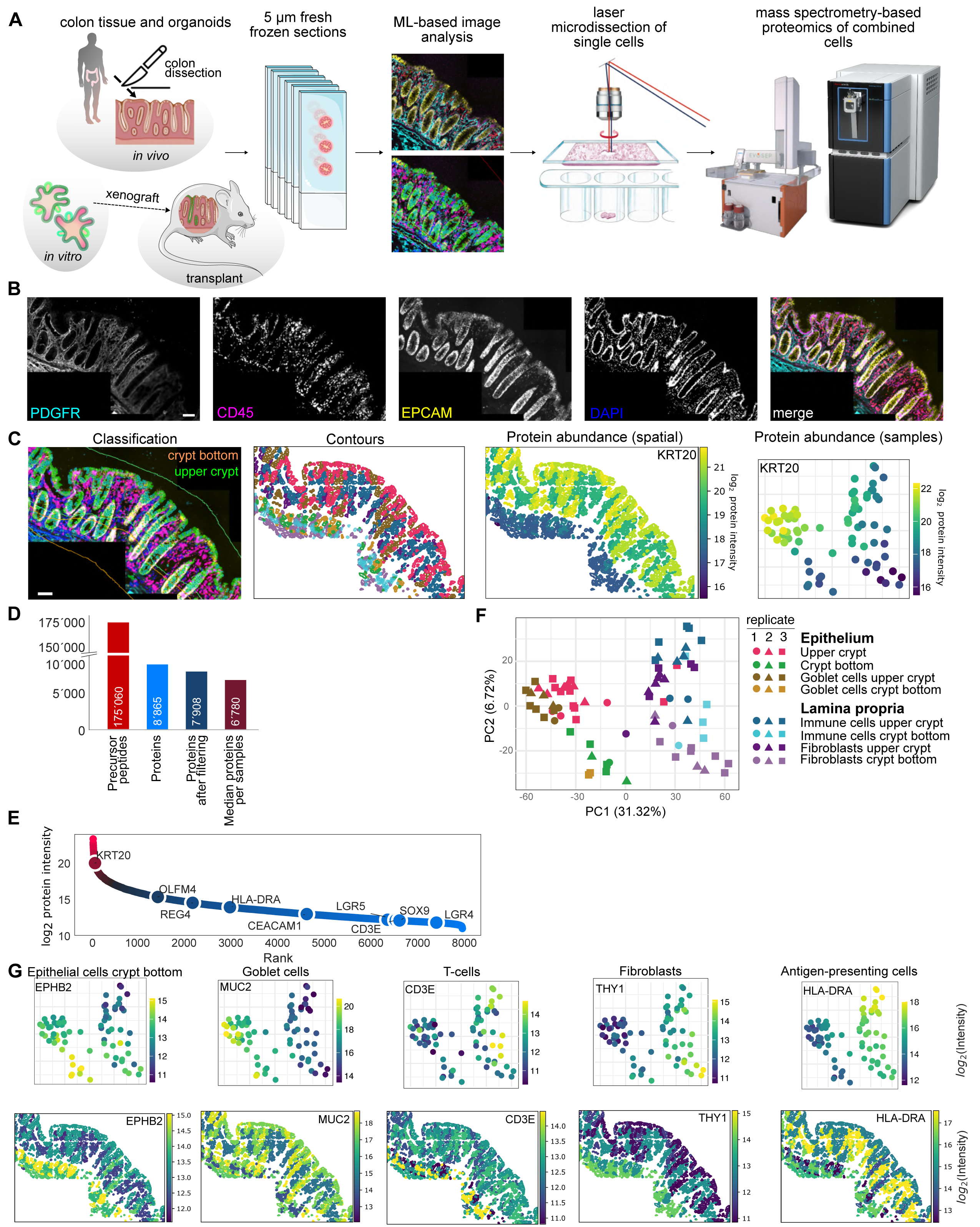
DVP analysis faithfully assesses cellular heterogeneity in the human colon. **A** Study design for the validation of organoids *in vitro* and organoid transplantation using Deep Visual Proteomics. **B** Immunofluorescence image of the human colon mucosa stained for fibroblasts (PDGFR), immune cells (CD45) and epithelial cells (EPCAM). **C** Crypt bottom and upper crypts were defined by a manually drawn line. Single cells were segmented and classified, contours exported, microdissected, and analyzed. This analysis reveals protein abundance across the colon mucosa and cell populations, as exemplified here for KRT20, a marker of differentiated epithelial cells. **D** Protein and precursor peptide identifications across all samples. **E** Median dynamic range of identified proteins across all samples after imputation and normalization. **F** PCA plot of samples isolated from the colon mucosa (three donors) as indicated by classification in C. **G** Protein abundance and spatial distribution of previously described cell type markers for different subpopulations in the human colon.

The analysis of the human colon mucosa included different populations of colonic epithelial cells (EPCAM^+^) and their microenvironment (lamina propria fibroblasts (PDGFRA^+^), immune cells (CD45^+^)) (Fig. 1B). Intestinal stem cells can be identified by *LGR5* expression^27^, but it has proven difficult to generate antibodies for reliable detection of LGR5. Alternative strategies for isolating human intestinal stem cells have been developed based on expression of EPHB2^28,29^, PTK7^30^ and OLFM4^31^, however, it remained challenging to detect epithelial stem cells in the human colon mucosa. We capitalized on the DVP technology to address this pertinent problem, enabling us to separate the epithelial crypt bottoms (enriched for stem cells, hereafter referred to as “crypt bottom”) from the upper part of the crypt (hereafter referred to as “upper crypt”) (Fig. 1C) based on spatial context. We used cellpose to segment high-resolution images for cell detection^32^. The resulting cell shapes and marker staining intensity were used to classify epithelial, goblet, immune cells and fibroblasts from the crypt bottom and upper crypt region using the biological image analysis software (BIAS) resulting in contours (one contour ≈ one cell in a 5 μm tissue section) (Fig. 1C). Technological limitations concerning availability of material and reliance on cellular markers for in-depth analysis of specific cellular subpopulations have so far hindered the characterization of functional states and phenotypes of human colonic epithelial cell subpopulations at protein levels. To address this, we isolated ∼500 contours per population by laser capture microdissection, lysed the collected contours, digested the proteins and performed proteome acquisition on the Evosep One liquid chromatography system coupled to an Orbitrap Astral mass spectrometer (Experimental Methods). With this approach, we achieved unprecedented sensitivity of cell populations directly isolated from fresh-frozen tissue, featuring 8,865 unique proteins across all cell populations and a median of 6,780 unique proteins per sample with a throughput of 40 samples per day (Fig. 1D-E, S1A-B). The limited sample amount from transplanted organoids restricted us to collecting a maximum of 100 contours from transplanted stem cells and ∼200 contours of transplanted epithelial cells in the upper crypt. Remarkably, the quantification of these samples still yielded ∼5,000 or ∼7,000 proteins, respectively (Fig. S1A).

Downstream principal component analysis (PCA) of the resulting data revealed that the samples from the human colon mucosa separated into two main clusters according to epithelium and lamina propria (immune cells and fibroblasts) along PC1, and further distributed according to the position along the crypt axis (bottom or top) along PC2 (Fig. 1F). To assess the reliability of identification and isolation of the different cell populations, we next assessed the abundance of previously described cellular markers for the isolated subpopulations in our sample set (Fig. 1C, G, S1C) and identified high expression of keratin (KRT)20 in upper crypt epithelial cells, Ephrin-type B receptor (EPHB)2 in crypt bottom epithelial cells, mucin (MUC)2 in goblet cells, thymocyte antigen (THY)1 in fibroblasts, as well as cluster of differentiation (CD)3E and human leukocyte antigen (HLA)-DRA in immune cells, thereby validating our human colonic mucosa proteome atlas. Interestingly, within the epithelial and lamina propria clusters, sample location along the crypt axis (upper crypt/bottom) rather than cell type drove their distribution (Fig. 1F). Differential activity of WNT and BMP signaling along the crypt axis regulate cellular organization, proliferation and differentiation within the intestinal epithelium, suggesting that these pathways might partially drive observed differences. The protein transgelin (TAGLN) was associated with the crypt bottom compartment irrespective of the cell type (Fig. S1D). In line with high WNT activity around the epithelial stem cell niche in the crypt bottom, TAGLN^+^ stromal cells have been identified as WNT producers^33^. The protein Zinc Finger ZZ-Type And EF-Hand Domain Containing (ZZEF)1, on the other hand, was enriched in the upper crypt compartment (Fig. S1D). ZZEF1 acts as a transcriptional regulator in cooperation with Krueppel-like factor (KLF)6 and KLF9^34^ which regulate IEC proliferation^35^ and absorption^36^, and might be modulated by the intestinal microbiota^37^, indicating a potential involvement in the integration of environmental stimuli into epithelial phenotypes. The interplay between luminal inputs and intrinsic regulation of mucosal gradients along the crypt axis and their molecular basis warrants further investigation.

In summary, we successfully generated a proteome atlas of the human colon mucosa in unprecedented depth with our DVP approach, which reveals differentially regulated protein levels along the crypt axis across cell types.

### DVP analysis reveals a robust correlation between human IECs *in vivo* and grown as organoids

For an in-depth characterization of human colon organoids at proteome level, we adapted the DVP pipeline described above to organoids. The accurate and sensitive assessment of functional cellular states at proteome level within a spatial context in combination with the *in vivo* proteome atlas as reference data set enables the benchmarking of model systems for human IECs (Fig. 1). Here we made use of a genetically engineered human colon cell organoid line, expressing the fluorescent reporter TdTomato under the control of the LGR5 promoter^12^ to identify epithelial stem cells (Fig. 2A). This allowed us to use the DVP workflow described above to identify, isolate and analyze human colonic stem cells (LGR5-TdTomato^+^ cells, hereafter referred to as “stem cells”), LGR5-TdTomato^-^ cells (hereafter referred to as “LGR5^-^ cells”), and goblet cells to generate a proteome atlas of human IECs grown as organoids *in vitro*. It should be noted that the half-life of the reporter protein might be longer than LGR5, thus TdTomato^+^ cells could contain a fraction of cells which have recently exited the stem cell state (e.g. transit amplifying progenitors). In the PCA, samples clustered according to different epithelial populations (Fig. 2B) with PC1 separating stem cells from the remaining IECs and PC5 separating goblet cells from stem cells and LGR5^-^ IECs. Expectedly, KRT20 was enriched in the LGR5^-^ cells (Fig. 2C), the stem cell marker EPHB2 in LGR5^+^ stem cells and MUC2 in goblet cells (Fig, 2C).

**Figure 2:**
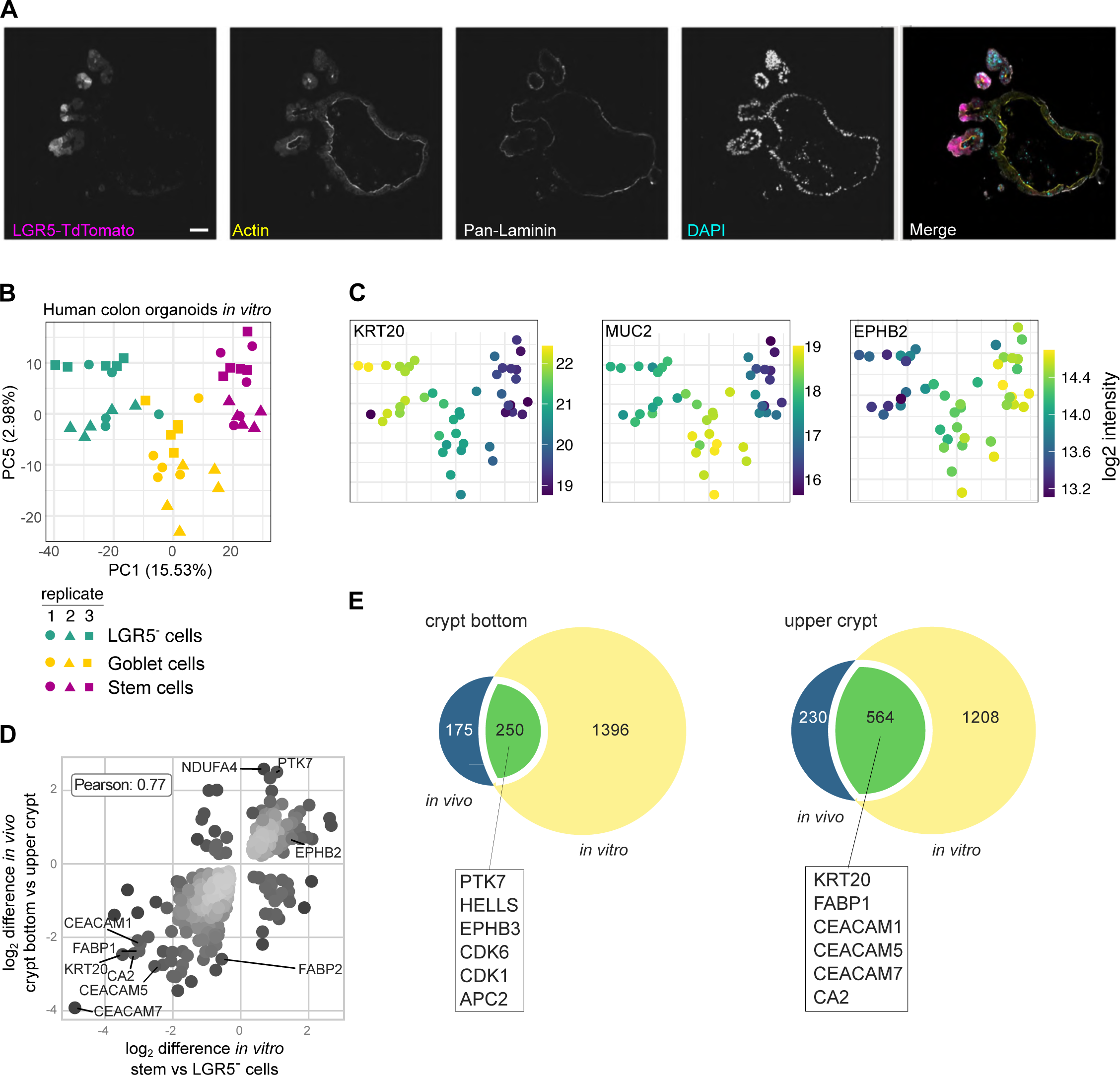
DVP analysis reveals a robust correlation between human IECs *in vivo* and grown as organoids. **A** Immunofluorescence image of a human colon organoid genetically engineered to express TdTomato under an LGR5 reporter for the identification of LGR5^+^ epithelial stem cells. **B** PCA plot of samples isolated from human colon organoids (three biological replicates (one organoid line, three separate passages), five technical replicates). **C** Abundance of previously described markers for different epithelial subpopulations (Krt20 – differentiated epithelial cells, MUC2 – goblet cells, EPHB” – stem cells). **D** Correlation plot of protein intensities of significantly changed proteins in epithelial cells located in the crypt bottom vs upper crypt *in vitro* and *in vivo*. **E** Venn Diagram of significantly changed proteins in epithelial cells in crypt bottom vs upper crypt *in vitro* and *in vivo*. Lines indicate selected overlapping proteins between *in vitro* and *in vivo* crypts.

A comparison of significantly changed proteins in stem versus LGR5^-^ cells measured *in vitro*, and those measured *in vivo* in the crypt bottom versus upper crypt respectively, showed a robust correlation between the lower and upper crypt compartments *in vivo* and *in vitro* (Pearson coefficient 0.77, Fig. 2D), which is in a similar range to the correlation of transcriptomes of murine small intestinal IECs *in vitro* and *in vivo* ^38,39^. Notably, ∼70% (crypt bottom) or ∼60% (upper crypt) of significantly enriched proteins in the respective populations *in vivo* were shared with organoids grown *in vitro* (Fig. 2E). Among these, we identified a number of described markers associated with the analyzed populations, indicating that key aspects of crypt bottom and upper crypt epithelial cells are preserved in *in vitro* culture. The higher number of proteins identified as differentially abundant *in vitro* is likely due to more homogenous populations isolated from *in vitro* than *in vivo* conditions (e.g., LGR5^+^ cells/crypt bottom), which allow for a more robust comparison.

To conclude, with our DVP approach we successfully benchmark human colon organoids to IECs *in vivo,* revealing a robust preservation of key compartment-associated features in organoids and highlighting their applicability as a model system for human colon IECs *in vitro*.

### Orthotopic transplantation reverts organoid phenotypes to an *in vivo-*like state

The transplantation of human organoids into the murine colon emerges as a novel model to dissect human IEC phenotypes and behavior within the mucosal environment^10,11^, but our current knowledge on how well human IECs transplanted into the murine colon recapitulate human IECs *in vivo* is limited to the assessment of selected markers for epithelial subpopulations^10,11^. To address this, we transplanted the genetically engineered human reporter organoids (Fig. 2A) into the murine colon (Fig. 3A). Consistent with previous reports, the cultured cells integrated into the murine colon mucosa and recapitulated the organotypic crypt structure featuring LGR5-TdTomato^+^ cells at the crypt bottom (Fig. 3B)^10–12^. For a comprehensive, unbiased assessment of epithelial phenotypes upon transplantation, we performed DVP analysis on the transplanted cells, focusing on stem (LGR5-TdTomato^+^, hereafter referred to as “stem cells”) and remaining cells (LGR5-TdTomato^-^, hereafter referred to as “LGR5^-^ cells”). Transplant size varies between mice and sometimes comprises only a few crypts. In protocols that require tissue dissociation (e.g. for scRNAseq), it can be challenging to efficiently recover these relatively rare cells. Furthermore, they often include lengthy enrichment steps such as cell sorting, which impacts IEC phenotypes^19^. For our DVP approach instead, we localized the transplants during sectioning, which enabled us to efficiently isolate transplanted IECs directly from their mucosal microenvironment. Strikingly, our DVP analysis revealed that transplanted organoids clustered with the *in vivo* IECs rather than organoid samples (Fig. 3C). This is particularly remarkable given that all organoid samples derive from the same organoid line (i.e., the same donor), while the IECs *in vivo* derive from three different donors, indicating that the phenotypic shift across conditions is stronger than interindividual differences.

**Figure 3:**
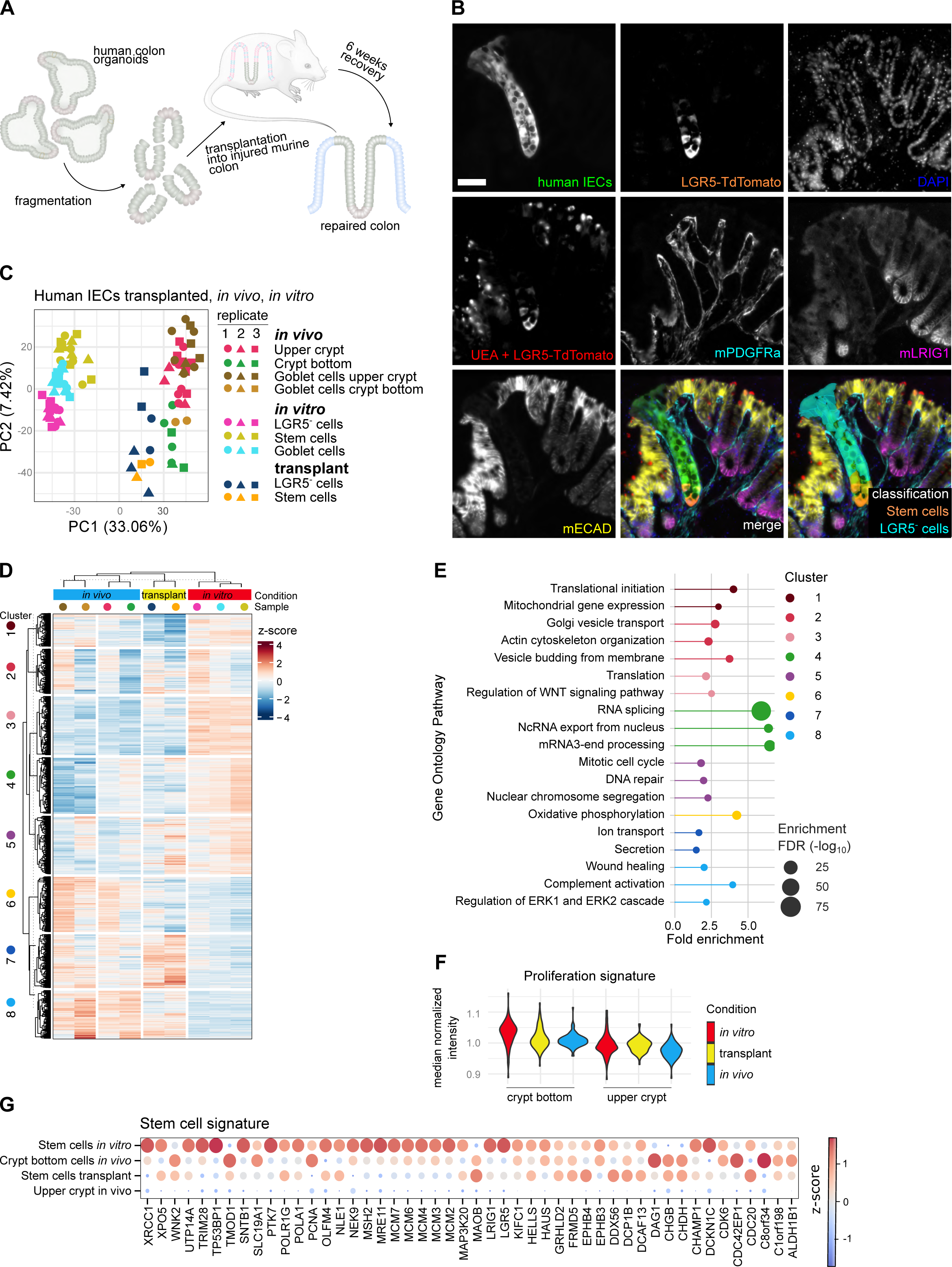
Human colon organoids transplanted into the murine colon recapitulate human colonocytes *in vivo*. **A** Workflow for orthotopic transplantation of organoids into the murine colon. **B** Immunofluorescence image human colon organoids (Fig. 2) transplanted into the murine colon (transplant). (GFP: human IECs. LGR5: stem cells (human). mECAD: epithelial cells (mouse). mPDGFR: fibroblasts (mouse). mLRIG1: crypt bottom compartment (mouse). UEA: mucus (goblet cells). **C** PCA plot of human colonocytes transplanted into the murine colon (one organoid line, three mice, one to three technical replicates), *in vitro* (organoids) and *in vivo* (human colon). **D** Heatmap of significantly changed proteins between organoids *in vitro*, transplanted organoids, and epithelial cells *in vivo*. **E** Gene ontology pathway enrichments of clustered proteins based on the heatmap in 3D. **F** Normalized protein intensities *in vitro*, in transplant, and *in vivo* of proteins that are associated with a proliferation signature in epithelial cells in the crypt bottom and the upper crypt. **G** Human stem cell proteome signature.

To gauge the biological magnitude of this shift, we included the lamina propria cells (fibroblasts, immune cells) isolated from the colon mucosa *in vivo* as outlier groups into the PCA (Fig. S3A). Surprisingly, despite the robust correlation between IECs *in vivo* and *in vitro* observed above, the distance between IECs grown as organoids *in vitro* and *in vivo* was very similar to the distance along PC1 between lamina propria cells and IECs *in vivo*, which are different cell types. A major driver for this differential clustering were components of the mucosal immunoglobulin A (IgA) (Fig. S3B), an important adaptive immune component of the mucosal barrier, which is secreted into the intestinal mucosa by B cells and subsequently transported into the intestinal lumen by IECs^40^. This indicates that the mucosal microenvironment has a significant impact on cellular proteomes across cell types, which should be considered when translating findings from organoid studies to *in vivo* phenotypes.

Collectively, the DVP analysis of orthotopically transplanted human colon organoids into the murine colon demonstrates that the cellular environment strongly impacts on IEC proteome profiles, pushing organoid phenotypes towards their *in vivo* counterparts.

To assess the cellular features driving phenotypic differences between IECs *in vitro* and within the mucosa, we performed a Kruskal-Wallis test across all epithelial samples. Hierarchical clustering of significantly changed proteins confirmed a separation of IECs grown *in vitro* from those isolated from the mucosa (*in vivo*, transplant) (Fig. 3D). Protein abundance patterns among these samples yielded eight clusters. Pathway analysis for the proteins within each cluster (Fig. 3E) revealed that signatures high in transcription (Cluster 4), translation (Cluster 1, 3), and proliferation (Cluster 5) characterized organoids cultured *in vitro* (partially shared with crypt bottom *in vivo* & transplanted stem cells), whereas *in vivo* and transplanted IECs were characterized by signatures associated with mucosal barrier function^41^ (e.g. complement activation, Cluster 8), functional features of mature IECs (e.g. ion transport, secretion, Cluster 7), and oxidative phosphorylation (Cluster 6). A direct comparison between IECs *in vivo* and *in vitro* confirmed these observations (Fig. S3C-H). The increased proliferative features *in vitro* were also evident as a specific enrichment of proteins involved in proliferation in IECs *in vitro*, which was decreased upon transplantation to levels similar to *in vivo* (Fig. 3F)^42^. To identify markers associated with upper crypt IEC phenotypes *in vivo*, we next assessed the PC loadings to identify proteins that drive the separation between IECs *in vivo* and *in vitro* (Fig. S3B). Here, carbonic anhydrase (CA)1 and MUC17 were amongst the highest scoring proteins. CA1 mediates ion transport, which is key for the regulation of water absorption in the intestine^43^. MUC17 is a membrane mucin forming the glycocalyx, an important barrier against bacterial attachment to the mucosa, which is compromised in IBD^44^ (Fig. S3I). In summary, components of two aspects of functional IECs *in vivo*, ion transport and barrier function, are underrepresented in IECs grown *in vitro* under the conditions tested here.

Altogether, our DVP approach revealed that the *in vitro* culturing conditions used here induce a high proliferation, low functional profile of IECs *in vitro*, and that these characteristics are reversible upon transplantation into the colon mucosa. This underscores the value of transplanted organoids as a system for the molecular dissection of epithelial phenotypes in a more *in vivo*-like setting, and highlights their applicability in regenerative medicine, e.g., for approaches to replenish impaired epithelium.

### Integrated DVP analysis identifies a human stem cell signature

The use of fluorescent reporters has enabled studies of intestinal epithelial stem cells in mice *in vivo* and in genetically engineered human organoids *in vitro* but it has so far been difficult to specifically isolate and analyze human stem cells *in vivo* due to the lack of antibody-stainable stem cell markers. Our study design uniquely allowed the collective in-depth proteome analysis of LGR5-TdTomato^+^ human stem cells *in vitro* and upon xenotransplantation in comparison to stem cell-enriched human IECs *in vivo*. The comparisons across these datasets enabled us to identify a shared protein profile enriched in stem cells *in vitro* and upon transplantation, and crypt bottom cells *in vivo*, which were downregulated in upper crypt cells *in vivo*. This human stem cell proteome signature includes 48 proteins (Fig. 3G) and as expected, contains a number of proteins associated with cell proliferation. The assessment of the expression patterns of these proteins via the Human Protein Atlas^45^ confirmed their localization at the crypt bottom *in vivo* (Fig. S4A-B). Notably, while all identified proteins localized within the stem cell niche, their abundance towards the crypt’s upper part varied (Fig. S4A-B). Based on this, we postulate that markers with a relatively confined expression such as EPHB3, meiotic recombination 11 (MRE11) and minichromosome maintenance complex component 2 (MCM2) could be suitable markers for a strongly stem cell-enriched IEC population. In comparison to previously published markers for stem cell enrichment in the human colon such as PTK7, EPHB2 and OLFM4 ^28,30,31^, expression of these markers is more restricted to the crypt bottom (Fig. S4B). EPHB3 is a receptor tyrosine kinase involved in regulation of stem cell positioning along the crypt axis and regulates mitogenic activity in cooperation with WNT^29,46^. As an antibody-stainable surface protein, we expect it to be a valuable marker for the enrichment of human stem cells, e.g. in cell sorting, which would address a major technical gap. MRE11^47^ and MCM2^48^ regulate DNA double-strand break repair and DNA replication, respectively. Other markers such as PCNA, MCM3, MCM4 likely include transit amplifying populations as well, in line with their roles in cell division^49–51^.

With this, our DVP approach has enabled the identification of EPHB3 as a potential novel surface marker for strong enrichment of stem cells, together with MRE11 and MCM2 as additional, antibody-stainable markers.

### WNT withdrawal induces upregulation of *in vivo* IEC markers

The protocols for expansion of IECs as organoids have been optimized for growth at the expense of differentiation. This is achieved via activation of the WNT pathway (supplementation of signals activating the canonical WNT pathway – WNT surrogate and R-spondin1), which is active in the crypt bottom compartment *in vivo*, and inhibition of BMP signaling (supplementation of Noggin), which is active in the upper crypt compartment *in vivo*^3,5,52^. We hypothesized that these conditions could be drivers of the observed *in vitro* characteristics shaped by high proliferation and lower functional features when compared to the *in vivo* and transplanted IECs. In line with this, both stem cells and LGR5^-^ cells *in vitro* were enriched for active WNT signaling^53^ when compared to their *in vivo* counterparts (Fig. S5A-B).

To address the impact of WNT and BMP signaling on epithelial phenotypes, we cultured organoids *in vitro* under conventional (+WNT, Noggin, RSPO (WNR)) or differentiation (-WNR +/- BMP) conditions^12,54^. We observed a clear shift in organoid proteome profiles upon withdrawal of WNR while the addition of BMP only had a minor additional effect (Fig 4A). As hypothesized, WNR withdrawal led to a decrease in WNT activation (Fig. S5C-D). It furthermore induced a downregulation of stem cell- and proliferation-associated proteins such as SOX9, MKI67, MCM2 and PCNA (Fig. 4B-C). This was also evident at a more global level when we assessed expression of proteins assigned to the proliferation signature^42^ and our stem cell signature identified above (Fig. 4D). At the same time, WNR withdrawal coincided with an upregulation of markers of mature IECs, such as KRT20, as well as CA1 and MUC17, which we identified in the analysis above as strongly associated with IECs *in vivo* (Fig. 4E). Similarly, the oxidative phosphorylation signature, which was enriched *in vivo* compared to organoids (Fig. S3F) was increased upon WNR withdrawal, indicating that IEC metabolic function is in part driven by IEC maturation state (Fig. 4F). Importantly, immunostaining of MUC17 in organoids upon WNR withdrawal revealed increased abundance of MUC17 at the apical surface, suggesting glycocalyx formation under these conditions. (Fig. 4G). Altogether, this indicates, as suggested previously, that withdrawal of WNR indeed drives organoids towards a more *in vivo*, upper crypt-like phenotype^55^.

**Figure 4:**
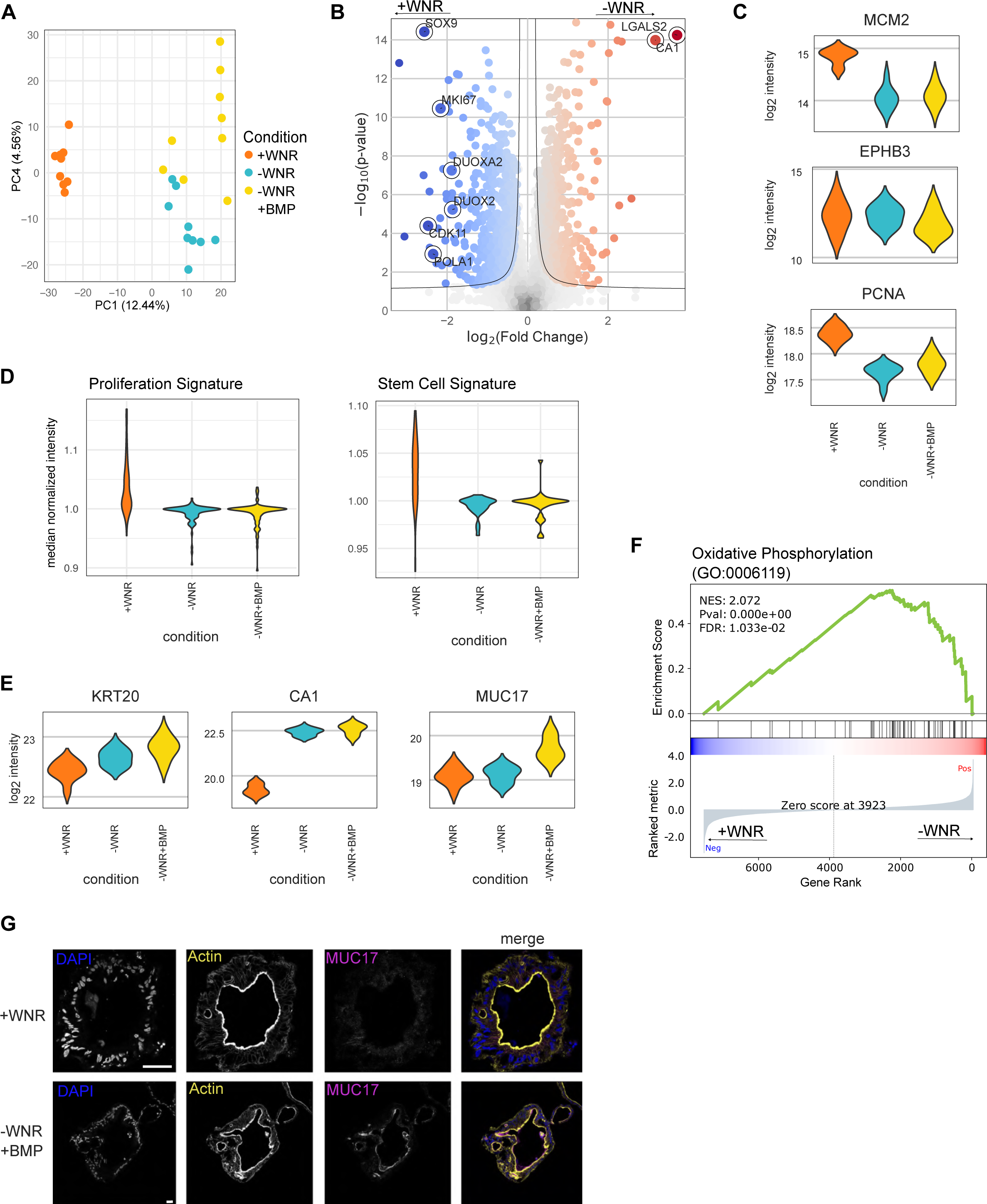
WNR withdrawal in colon organoids cultured *in vitro* induces upregulation of *in vivo* IEC markers. **A** PCA of organoids cultured with WNR (WNT3a (W), Noggin (N), R-spondin-3 (R))(+WNR), without WNR (-WNR), and with BMP (Bone Morphogenetic Protein) but without WNR (-WNR +BMP). **B** Volcano plot of organoids cultured with WNR and without WNR. **C** Decrease of stem cell markers of colonic epithelial cells by withdrawal of WNR and addition of BMP. **D** Median normalized intensity of a proliferation signature^41^ and stem cell signature in +WNR, -WNR, and -WNR +BMP. **E** Increase of differentiation markers of colonic epithelial cells by withdrawal of WNR and addition of BMP. **F** Fluorescence microscopy showing the increase of MUC17 in colon organoids upon withdrawal of WNR and addition of BMP. **G** Gene Set Enrichment Analysis showing an increase of the oxidative phosphorylation Gene Ontology pathway in -WNR vs +WNR.

## Discussion

We here employ DVP to generate an in-depth proteome atlas of the human colon mucosa, which we use to benchmark human colon organoids grown *in vitro* and upon orthotopic xenotransplantation. We originally developed DVP as a spatial proteomics technology that enabled the acquisition of the proteome of about 10 samples per day, quantifying up to 5,000 proteins from input material equivalent to 100 – 200 cells^25^. In our improved workflow, which includes coupling the Evosep One liquid chromatography system to the Orbitrap Astral analyzer, throughput is increased to 40 samples per day. Remarkably, total proteome acquisition time for this in depth, functional organoid study encompassing 136 samples was only 88 hours. Despite faster acquisition, we increased the proteome depth to a total of 8,865 unique proteins. This setup also enabled the quantification of ∼5,000 proteins from as little as 100 transplanted stem cell contours, corresponding to only 20 intact cells. The increased proteome depth was essential to enable conducting this study since it enabled us to identify low abundant proteins such as SOX9 or LGR5 from cells dispersed over several slides.

Based on this improved DVP pipeline, the benchmarking of human colon organoids reveals a robust correlation of IECs grown *in vitro* and *in vivo*. Nevertheless, IECs grown *in vitro* display high proliferation and altered functional and metabolic signatures compared to *in vivo*, which has important implications for the use of organoids as models to dissect epithelial phenotypes. We show that these features are driven by organoid culture conditions and are largely reverted upon organoid transplantation into the murine mucosa, as well as, in part, by altering organoid culturing conditions (WNR withdrawal). Altogether, our study validates the applicability of orthotopically xenotransplanted organoids as tools to mechanistically dissect human IEC phenotypes in an *in vivo*-like setting and highlights their potential to accurately replenish the intestinal epithelium in a regenerative medicine approaches.

Human organoid models are instrumental for assessing key biological questions in a human context. The premise that the organoid model truly recapitulates *in vivo* phenotypes, and an awareness of its limitations, is crucial for the translatability of *in vitro* results to *in vivo* applications. A key gap currently limiting the exploitation of the full potential of human organoids in biomedical research is the characterization and validation of organoids as accurate models for human biology^1,7,13^. An in-depth characterization of native IEC states within their *in vivo* environment is essential to establish a reference for benchmarking of human-like model systems. We have here tackled this issue, using our DVP approach to generate an in-depth proteome atlas of the homeostatic human colon, which serves as an important reference for future studies assessing e.g. disease-associated changes in the human colon. Notably, the DVP setup does not require fresh tissue dissociation and enrichment of living cells, which reduces the impact of lengthy isolation protocols on cellular phenotypes and thereby enabled us to assess the proteomes of mucosal cell types in their native state. We successfully identified and differentiated the isolated mucosal cell populations. Interestingly, aside from cell type-specific protein abundance patterns, we observed location-skewed protein abundance along the mucosal crypt axis. A similar zonation has been reported previously for murine small intestinal epithelial cells at transcriptome level^14^, and it is well known that differences in e.g. WNT and BMP signaling along the crypt axis regulate epithelial phenotypes^1^. We here address this comprehensively across the different cell types in the mucosa at the protein level and identify the proteins ZZEF1 and TAGLN, which associate with the upper or crypt bottom compartment across the analyzed cell types, respectively. In the future, it will be interesting to study this protein regulation along the crypt axis in further detail and to dissect how e.g. WNT and BMP signaling gradients, as well as luminal cues such as microbiota shape protein abundance and cellular identity. This will shed light on the regulatory pathways maintaining tissue structures which are key for intestinal homeostasis and abrogated for example in the context of colorectal cancer^56,57^.

Our human colon proteome atlas further enabled us to benchmark widely used *in vitro* ^3–5^ and emerging organoid transplantation models^10^ for human IECs. Importantly, while we detect robust proteome correlation between IECs grown *in vitro* and their *in vivo* counterparts, which mirrors previous reports on transcriptome level in murine small intestine, we observe a striking phenotypic switch of organoids upon transplantation into the mucosa, rendering them *in vivo*-like. A major difference between organoids grown *in vitro* and transplanted into the mouse colon is a reduction in the proliferation signature, comparable to *in vivo* IECs, upon reintroduction into the mucosa. In addition to the high proliferation state, organoids grown *in vitro* display lower functional features (e.g. ion transport), as well as a different metabolic signature characterized by lower oxidative phosphorylation. In the murine small intestine, oxidative phosphorylation has been linked to the regulation of stem cell identity and differentiation into Paneth cells^58^. We here find that proteins associated with oxidative phosphorylation are, at least in part, differentially regulated depending on epithelial maturation state. It remains to be shown whether this correlates with actual changes in metabolism between epithelial subpopulations, and whether/how epithelial differentiation and metabolism are linked in colonic IECs^59^. We further make use of our dataset to identify CA1 (ion transport/water homeostasis^43^) and MUC17 (glycocalyx in the brush border of differentiated IECs/barrier function^44^) as markers for human upper crypt IECs *in vivo*.

We show that high proliferation and low functional features observed in IECs grown *in vitro* are driven by the culture conditions (high WNT, low BMP signaling), rather than an intrinsic cellular feature selected for during culture, and that this state, including abundance of CA1 and MUC17, can be partially reverted by adjustments in culturing conditions (-WNR +BMP). Notably, recent advances in organoid-on-a-chip models using hydrogels which recapitulate the mucosal crypt structure and molecular gradients, feature similar IEC shifts to a more *in vivo*-like phenotype at transcriptome level^60^. These findings have important implications for the use of organoids to study IEC functions *in vitro*, especially when focusing on the role of upper crypt IECs, e.g. in host-microbe interactions.

The phenotypic reversion of organoids transplanted into the murine colon to a more *in vivo*-like phenotype highlights a remarkable homology between mouse and human stem cell niche factors. A more detailed analysis of the differences between transplanted organoids and IECs *in vivo* will reveal which molecular pathways drive the difference we observed between these two populations. One key aspect aside from the limited compatibility of mouse and human growth factor signaling could be the fact that we used immunocompromised mice for the xenotransplantation to prevent rejection. Future studies comparing human to murine organoids transplanted into the murine colon will be able to dissect the impact of species-specificity and the presence of immune cells on transplanted epithelial cells.

Finally, we capitalize on the unprecedented possibility to characterize human LGR5^+^ stem cells in the colon mucosa to identify a human stem cell proteome signature, which reveals EPHB3, MRE11 and MCM2 as antibody-stainable markers for the enrichment of human colonic stem cells *in vivo*. Notably, expression of these markers is more strongly restricted to the crypt bottom *in vivo* compared to previously published markers for the enrichment of human stem cells (EPHB2, PTK7, OLFM4). As EPHB3 is a surface protein, we expect that this marker will be of great value for the community to identify and isolate stem cell-enriched IECs for future studies of human intestinal stem cells. Furthermore, this showcases the strength of DVP to i) efficiently isolate rare cells from tissue in their native state and to ii) use proteome data to directly identify antibody-stainable markers. In addition, it serves as a proof-of-principle for the specific isolation and analysis of genetically modified xenotransplanted human IECs from the murine colon and lays the base for future mechanistic studies, e.g. in the context of tissue damage and repair, and host-microbe interactions.

We here advanced the DVP pipeline, demonstrating that DVP is a uniquely well-suited methodology for the faithful in-depth analysis of functional cellular phenotypes in a densely packed tissue like the colon mucosa. Limited sensitivity has so far been a major difficulty for the use of proteomics to dissect dynamic tissue processes, especially in the context of tightly regulated responses such as inflammation (i.e., low abundant, spatially restricted proteins). An additional limitation has been the ability to isolate cells in a near to native state, in the absence of alterations by tissue handling including single cell isolation. The DVP protocol we use here tackles these hurdles, enabling higher throughput and requiring less input material than the original method, and preserving spatial context while reducing the impact of isolation protocols on cellular phenotypes. These technological advancements are promising regarding the expansion of DVP for the acquisition of proteomes of single cells^61^. This opens exciting perspectives for the use of DVP to study dynamic tissue processes such as inflammation, even from rare patient material.

Taken together, the presented data has important implications for the selection of *in vitro* organoid systems to study specific aspects of epithelial cell biology. The phenotypic reversion of organoids transplanted into the murine colon to a more *in vivo*-like phenotype highlights the impressive homology between mouse and human stem cell niche factors, underlines the suitability of the murine (orthotopic transplantation) model for studies of epithelial-niche interactions with a translational perspective and opens exciting possibilities for the use of organoid transplantation in regenerative medicine.

## Methods

### Human colon mucosa samples

All individuals included in this study were attending the Department of Gastroenterology, Herlev Hospital, University of Copenhagen, Denmark, for the Danish National Screening Program for Colorectal Cancer or evaluated for various gastrointestinal symptoms but were included only if all subsequent examinations were normal. The exclusion criteria included age below 18 or over 80 years; impaired cognitive functions, e.g., dementia; pregnant or lactation women; ongoing treatment with anticoagulation, and patients unable to understand Danish language. The study was approved by the Scientific Ethics Committee of the Capital Region of Denmark (reg. no. H-21038375). All individuals were informed of the study both orally and in writing, in compliance with the Declaration of Helsinki and the guidelines of the Danish National Scientific Ethics Committee. Written informed consent was obtained prior to inclusion.

For those individuals included, human colon mucosa samples (cancer-associated bowel resection or biopsies (healthy individuals undergoing cancer screening)) were immediately transferred to 4% PFA (Sigma) upon sampling and fixed at 4 °C for 2-10 days, depending on sample size. Samples were then washed in PBS and transferred to 30% sucrose/PBS and dehydrated for 2-10 days at 4 °C. Next, samples were embedded in OCT, frozen on dry ice and stored at −80C until further analysis.

### Human colon organoid culture

Human colon organoids were cultured as previously described^5^. Briefly, upon single cell dissociation, 3,000 – 4,000 single cells were seeded in 30 µL Matrigel domes and maintained in advanced DMEM/F-12, supplemented with penicillin-streptomycin, 10 mM HEPES, 2 mM GlutaMAX, 100 ng/mL recombinant mouse Noggin, 1x B27, 500 nM A83-01, 1% NGS-WNT, 1 mg/mL recombinant human R-spondin-1, 100 ng/mL recombinant human IGF, 50 ng/mL recombinant human FGF2, 1 mM N-Acetylcysteine and 10 nM recombinant human Gastrin. For WNR withdrawal, organoids were cultured in conventional medium until d7. Organoids were then reseeded in fresh Matrigel domes (no splitting) and maintained until d10 in advanced DMEM/F-12, supplemented with penicillin-streptomycin (Penstrep), 10 mM HEPES, 2 mM GlutaMAX, 1x B27, 500 nM A83-01, 100 ng/mL recombinant human IGF, 50 ng/mL recombinant human FGF2, 1 mM N-Acetylcysteine and 10 nM recombinant human Gastrin in the presence or absence of BMP4 (10 ng/ml). Organoids were split every 7d for maintenance. Organoids were harvested at d10 for the analyses presented in this study. Human colon organoids from healthy individuals have been used for this study. The LGR5-TdTomato reporter organoid line has been described before^12^. To introduce a constitutive GFP reporter to the cells for easier localization of the transplant, eight wells (i.e. eight 30 µL Matrigel domes) of organoids were mechanically disrupted, washed and resuspended ∼600 µL media supplemented with Y-27632 (10 uM). Lenti virus was added to the cells to transduce them with a plasmid expressing GFP under the SFFV promotor^62^. The cells were incubated for 4 h at 37 °C, washed three times in DMEM medium and subsequently seeded into four Matrigel domes (30 µL). After three days of culture, transduced cells were selected by addition of 2 ug/ml Puromycin to the media. Cells were passaged twice, tested according to FELASA standards (IDEXX), and subsequently used for transplantation.

For cryosamples, 500 µL ice cold cell recovery solution was added to each well. Matrigel domes were carefully scraped off with a cut open P1000 pipet tip and transferred to 5 ml cell recovery solution (R&D systems) on ice. After 30 min, the supernatant was removed, organoids were resuspended in 4% PFA and fixed for 1h at ambient temperature. Subsequently, organoids were washed three times in 5 ml PBS (if necessary, organoids were spun down for 2 min at 100g), embedded in OCT (Tissue Tek) in cryomolds, and frozen on dry ice. Samples were stored at −80 °C until further analysis.

For bulk proteome analysis, organoids were harvested as previously described^63^. Briefly, 1 ml ice cold 0.1% BSA/PBS was added to each well and matrigel domes were broken up by pipetting 10 times with a P1000 pipet. Organoids from four wells were pooled per sample in a tube containing 3 ml 0.1% BSA/PBS. Cells were pelleted by centrifugation (5 min, 300 x *g*, 4 °C), supernatant was removed and cells were resuspended in 1 ml 0.1% BSA/PBS and pelleted again. Upon removal of the supernatant, cells were resuspended in 200 µL 0.1% BSA/PBS and transferred to a 1.5 ml Eppendorf tube (pre-coated with 0.1% BSA/PBS) and kept on ice until further processing.

### Orthotopic xenotransplantation

NOD.Cg-Prkdc^scid^ Il2rg^tm1Sug^/JicTac (NOG) mice were used for transplantation assays. All animal procedures were approved by the Danish Animal Inspectorate (license number 2018-15-0201-01569 to Kim B. Jensen).

In preparation of the transplantation, organoids were grown as described above until d5-6 in 6-well plates containing nine Matrigel domes per well. 3 ml ice cold cell recovery solution was added to each well. Matrigel domes were carefully scraped off with a cut open P1000 pipet tip and transferred to 5 ml cell recovery solution (R&D systems) on ice for 20 min. Cells were subsequently pelleted for 3 min at 300 x *g*, washed once in PBS and resuspended in 200 µL of 5% Matrigel/PBS per mouse. Right before transplantation, organoids were dissociated by pipetting 20x with a pre-wet P1000 pipette.

Transplantation was performed as described previously^11^, with slight modifications. Mice were anesthetized with 2% isoflurane before the procedure. The colon content was flushed with PBS and an electric interdental brush, soaked in prewarmed 0.5 M EDTA, was used to brush crypts off on one side of the colon. The organoids suspension was subsequently infused into the conditioned colon. Glue (Histo-acryl, B. Braun) was added to the anal verge and left for 3h to avoid the ejection of the organoid suspension and thereby enhance the engraftment of the infused material. Mice were monitored daily. Transplanted samples were isolated six weeks after transplantation. For cryosectioning, the colon was isolated, cut open and placed under a fluorescent microscope (Evos) to locate GFP^+^ transplanted cells. The colon area containing the transplant was subsequently cut out, fixed in 4% PFA at 4 °C over night, dehydrated in 30% sucrose/PBS over night at 4 °C and then embedded in OCT and frozen on dry ice. Samples were kept at −80 °C until further analysis.

### Cryosectioning, immunofluorescent staining and imaging for DVP

2-mm-thick polyethylene naphthalate membrane slides (Zeiss) were pretreated by ultraviolet ionization for 3 h. Without delay, slides were consecutively washed for 5 min each in 350 ml acetone and 7 ml VECTABOND reagent to 350 ml with acetone, and then washed in ultrapure water for 30 s before drying in a gentle nitrogen air flow. The slides were treated with a dilution of 7 mL Vectabond in 350 mL acetone for 5 minutes without prior washing in acetone or subsequent washing in water. Afterwards, the slides were dried in an incubator at 30 °C for 3 hours.

Frozen samples in OCT were cut with a Leica cryostat in 5 um sections. Samples were subsequently dried for 1h at ambient temperature, rehydrated with 500 µL PBS for 1 min and permeabilized with 300 ul PBS/0.5% TritonX-100. Tissue sections were blocked in 200 µL PBS/donkey serum for 30 min at room temperature and subsequently incubated with the primary antibody mix in blocking buffer overnight at 4C. The next day, samples were washed three times with 500 µL PBS and incubated for 40 min at ambient temperature with the secondary antibody mix in PBS. Upon washing three times with PBS, samples were mounted using anti-fade fluorescence mounting medium (abcam). Samples were subsequently imaged as described below and, if necessary subjected to a second round of staining. For this, samples were bleached using bleaching buffer (24 mM NaOH and 4.5% H2O2) for 10 min at room temperature, washed with PBS and stained as above.

Antibodies and staining reagents used in this study: CD45-BV421 (30-F11, Biolegend, 1:100), Lrig1 (R&D Systems AF3688, 1:50), PDGFR (EPR22059-270, abcam, 1:100), UEA-Atto550 (Atto-Tec, 1:500), EPCAM-APC (EBA1, BD Biosciences, 1:50), EPCAM-APC (G8.8, Fisher Scientific,1:50), ECAD (ECCD2, Thermo Fisher, 1:200), CD45 (HI30, Stem cell, 1:200), DAPI (Sigma), MUC17 (Merck HPA031634, 1:200), CA1 (EPR5193, abcam, 1:200), Pan-Laminin-AF647 (Novus Biologicals NB300-144AF647, 1:100).

The samples were imaged on a Zeiss Axioscan 7 microscope slide scanner at a magnification of 20×, with three z-layers with intervals of 2.5 mm. Human colon tissues were imaged in two secutive rounds. For the first round, the acquisition settings were 4 ms illumination time and 1.49% 385 nm laser for DAPI, 20 ms illumination time and 100% 475 nm laser for AF488, and 300 ms illumination time and 100% 735 nm laser for AF750. For the second round, the acquisition settings were 4 ms illumination time and 1.49% 385 nm laser for DAPI, 15 ms illumination time and 100% 475 nm laser for AF488, 60 ms illumination time and 100% 567 nm laser for AF568, and 20 ms illumination time and 100% 630 nm laser for AF647. For *in vitro* organoids, the acquisition settings were 2 ms illumination time and 1.1% 385 nm laser for DAPI, 2.2 ms illumination time and 100% 475 nm laser for FITC, 30 ms illumination time and 100% 567 nm laser for Rhoda, and 8 ms illumination time and 100% 630nm laser for AF647. Transplanted organoids were imaged in two staining rounds. The first round was imaged with an illumination time of 1.2ms and 1.5% 385 nm laser for DAPI, 3 ms illumination time and 100% 475 nm laser for Af488, 80 ms illumination time and 100% 567 nm laser for tdTomato, 20 ms illumination time and 100% 630 nm laser for Af647, and 100 ms illumination time and 100% 735 nm laser for Af750.

### Image Analysis

Corresponding images of the two imaging rounds were cropped and subsequently concatenated in imagej. Afterwards, the images were registered using the RigidBody transformation in HyperStackReg on the GFP and tdTomato channel in the transplanted organoids and DAPI in the *in vivo* human colon, and all channels were merged.

Images were split into tiles using the Biological Image Analysis Software (BIAS, Single-Cell Technologies Ltd.) and each tile was segmented in Napari using the cellpose cytosolic algorithm in the serialcellpose plugin. Images were not treated as RGB, batch size was set to 3, flow threshold was set to 3, cell probability threshold was set to –4, diameter was set to 30, the magenta channel was set as channel to segment, and the yellow channel was used as a helper channel. Image analysis was continued in in BIAS by filtering shapes for a minimum size of 50 µm^2^ and a maximum size of 2000 µm^2^. Features of segmented cells were extracted and classified using a multi-layer perceptron classifier with default settings. For human colon tissue, the bottom part of crypts was manually annotated using the region feature to distinguish stem cells and differentiated epithelium. Contours of cells were sorted using the “Greedy” setting and coordinates of the contours were exported.

### Laser Microdissection

Contours were imported at 63× magnification, and laser microdissection performed with the LMD7 (Leica) in a semi-automated manner at the following settings: power 46, aperture 1, speed 40, middle pulse count 4, final pulse 8, head current 46-50%, and pulse frequency 2,600. Contours were sorted into a low-binding 384-well plate (Eppendorf 0030129547). 500 contours were collected per sample except for immune cells surrounding upper crypt of which 700 contours were collected. Due to limited sample amount in the transplanted organoids, 200 contours were collected for differentiated cells and about 100 contours were collected for stem cells. An overview of collected biological replicates and technical replicates per cell population can be found in the supplementary data (Table S1). Contours were rinsed to the bottom of the well by filling the wells up with 40 mL acetonitrile, vortexing for 10 seconds, and centrifuging at 2000 x *g* at ambient temperature for 5 min. A SpeedVac was used to evaporate the acetonitrile at 60 °C for 20 min or until achieving complete dryness and the contours were stored at 4 °C.

### DVP proteome sample preparation and acquisition

Lysis was performed in 4 mL of 0.01 % n-dodecyl-beta-maltoside in 60 mM triethyl ammonium bicarbonate (TEAB, pH 8.5, Sigma) at 95 °C in a PCR cycler with a lid temperature of 110 °C for 1 h. 1 mL of 60% acetonitrile in 60 mM TEAB was added and lysis continued at 75 °C for 1 h. Proteins were first digested with 4 ng LysC at 37 °C for 3 h and subsequently digested overnight using 6 ng trypsin at 37 °C. The digestion was terminated by adding 1.5 mL 5 % TFA. Samples were dried in a SpeedVac at 60 °C for 40 min and stored at −80 °C.

C-18 tips (Evotip Pure, EvoSep) were washed with 100 µL of buffer B (0.1% formic acid in acetonitrile), activated for 1 min in 1-propanol, and washed once with 20 µL buffer A (0.1% formic acid). Samples were resuspended in 20 mL buffer A on a thermoshaker at room temperature at 700 x *g* for 15 min. Peptides were loaded on the C-18 tips, washed with 20 mL buffer A, and then toped up with 100 mL buffer A. All centrifugation steps were performed at 700 x *g* for 1 min, except peptide loading at 800 x *g* for 1 min.

Samples were measured with the Evosep One LC system (EvoSep) coupled to an Orbitrap Astral mass spectrometer (Thermo Fisher). Peptides were separated on an Aurora Elite column (15 cm x 75 mm ID with 1.7 mm media, IonOpticks) at 40 °C running the Whisper40 gradient. The mobile phases were 0.1% formic acid in liquid chromatography (LC)–MS-grade water (buffer A) and 0.1% formic acid in acetonitrile (buffer B). For samples consisting of 500 contours, the Orbitrap Astral MS was operated at a full MS resolution of 240,000 with a full scan range of 380 − 980 m/z. The AGC target was set to 500% for full scans and fragment ion scans. Fragment ion scans were recorded with a maximum injection time of 5 ms and with 300 windows of 2 Th scanning from 150 − 2000 m/z. Fragmentation of precursor ions took place using HCD with 25% NCE. Samples consisting of 200 contours (stem cells from transplanted organoids) were acquired using a full maximum injection time of 100 ms for MS1. Fragment ion scans were recorded with a maximum injection time of 14 ms (MS2), an AGC target of 800 %, and with 75 windows of 8 Th scanning from 150 − 2000 m/z.

### DVP raw MS data analysis

Raw files were converted to mzML using MSconvert and analyzed in DIA-NN 1.8.1 using an in-silico DIA-NN predicted spectral library (101370 protein isoforms, 177027 protein groups and 7821224 precursors in 3872218 elution groups)^64^. A human proteome reference database, including isoform information and the tdTomato fluorophore sequence, was used to generate the library and search the raw files (Uniprot March 2023). Following configuration was set for the search: N-terminal methionine excision was enabled, digest was performed at K* and R*, maximum number of missed cleavages was set to 2, maximum number of variable modifications was set to 2, oxidation of methionine was considered as variable, acetylation of the N-terminus was considered as variable, Protein inference = “Genes”, Neural network classifier = “Single-pass mode”, Quantification strategy = “Robust LC(high precision)”, Cross-run normalization = “RT-dependent”, Library Generation = “Smart profiling”, and Speed and RAM usage = “Optimal results”. Mass accuracy and MS1 accuracy were set to 15. “Use isotopologues”, “No shared spectra”, “Heuristic protein inference” and “MBR” were activated.

### DVP data analysis

Data analysis was mostly performed in Perseus and AlphaPeptStats ^65,66^. Python and R were used to conduct further analyses and visualize the data. The first technical replicate of the second biological replicate of fibroblasts at the bottom of crypts (fib_top_02_01) was removed due to the quantification of less than 2000 proteins. Raw data was imported into Perseus, and proteins filtered for 80 % data completeness within samples of the same cell type and same location in the human tissue. Missing values were replaced from a normal distribution with a width of 0.3 and a down shift of 1.3. Data was normalized by aligning the median intensity of all samples. Median intensities of each sample were determined, and the median of these median intensities was divided by the median of each sample. The resulting factor was multiplied with each intensity of the sample. Differential abundance analyses for volcano plots and enrichment analyses were performed in Perseus. Kruskal-Wallis tests were performed in Perseus with Benjamini-Hochberg FDR correction and a threshold of 0.01. GSEAs were performed using the GSEApy (v 1.0.6) package against the GO_Biological_Process_2023 dataset^67,68^.

### Bulk Proteome sample preparation and acquisition

200 mL 60 mM TEAB lysis buffer was added to the washed and pelleted organoids. Samples were lysed at 95 °C shaking at 800 rpm for 30 min. Afterwards the lysate was sonicated at 4 °C in 30 s intervals for 10 min. 18 mL ACN was added to bring the lysis buffer to a final concentration of 12.5 % ACN and lysis continued at 95 °C shaking at 800 rpm for another 30 min. Debris was pelleted at 4 °C at 20,000 x g for 10 min and supernatants transferred to fresh tubes. Protein concentration of supernatants was determined using nanodrop and 200 mg were used for further processing. Lys-C and trypsin were added at a protein to enzyme ratio of 50:1. Digestion took place at 37 °C shaking at 800 rpm overnight. Peptides were lyophilized using a SpeedVac at 60 °C for 1 hour. Peptides were resuspended in 200 mL Evosep buffer A (0.1 % formic acid) and 60 mL corresponding to 60 mg were loaded in triplicates on 3 layers of SDB-RPS membranes. About 10 ng were loaded on Evotips Pure.

Samples were measured with the Evosep One LC system (EvoSep) coupled to an Orbitrap Astral mass spectrometer (Thermo Fisher). Peptides were separated on an Aurora Elite column (15 cm x 75 mm ID with 1.7 mm media, IonOpticks) at 40 °C running the Whisper40 gradient. The mobile phases were 0.1% formic acid in liquid chromatography (LC)–MS-grade water (buffer A) and 0.1% formic acid in acetonitrile (buffer B). The Orbitrap Astral MS was operated at a full MS resolution of 240,000 with a full scan range of 380 − 980 m/z and a maximum injection time of 100 ms. The AGC target was set to 500% for full scans and fragment ion scans. Fragment ion scans were recorded with a maximum injection time of 5 ms and with 300 windows of 2 Th scanning from 150 − 2000 m/z. Fragmentation of precursor ions took place using HCD with 25% NCE.

### Bulk proteome raw MS data analysis

Raw files were converted to mzML using MSconvert and analyzed together with the DVP samples in DIA-NN 1.8.1 using an in-silico DIA-NN predicted spectral library (101370 protein isoforms, 177027 protein groups and 7821224 precursors in 3872218 elution groups)^64,69^. A human proteome reference database, including isoform information and the tdTomato fluorophore sequence, was used to generate the library and search the raw files (Uniprot March 2023). Following configuration was set for the search: N-terminal methionine excision was enabled, digest was performed at K* and R*, maximum number of missed cleavages was set to 2, maximum number of variable modifications was set to 2, oxidation of methionine was considered as variable, acetylation of the N-terminus was considered as variable, Protein inference = “Genes”, Neural network classifier = “Single-pass mode”, Quantification strategy = “Robust LC(high precision)”, Cross-run normalization = “RT-dependent”, Library Generation = “Smart profiling”, and Speed and RAM usage = “Optimal results”. Mass accuracy and MS1 accuracy were set to 15. “Use isotopologues”, “No shared spectra”, “Heuristic protein inference” and “MBR” were activated.

### Bulk proteome data analysis

Data analysis was mostly performed in Perseus and AlphaPeptStats. Python and R were used to conduct further analyses and visualize the data. Raw data was imported into Perseus, and proteins filtered for 80 % data completeness within samples of the same cell type and same location in the human tissue. Missing values were replaced from a normal distribution with a width of 0.3 and a down shift of 1.3. Differential abundance analyses for volcano plots and enrichment analyses were performed in Perseus and visualized in python and R. GSEAs were performed using the GSEApy (v 1.0.6) package against the GO_Biological_Process_2023 dataset.

## Author contributions

Conceptualization: FP, AH, AM, KJB, MM. Experimentation: FP, AH, SK, SS, AB, DLC, JS. Reagents and material: TLB, TS, CS, OHN. Writing – original draft: FP, AH, KJB, MM. Writing – review and editing: all authors.

## Acknowledgements

The authors thank Daniela Mayer and Hjalte L. Larsen for critical reading of the manuscript, Kira Petzold for preparing pretreated membrane slides, Xiang Zheng for providing staining protocols, and the Mann and Jensen groups for fruitful discussions. We acknowledge support by the CPR/reNEW imaging facility as well as Core Facility for Microscopy. This work was funded by grants from the Novo Nordisk Foundation (NNF14CC0001, NNF15CC0001 to MM and NNF18OC0034066, NNF20OC0064376 to KBJ)) and the Independent Research Fund Denmark (0134-00111B) to KBJ. Additionally, SK and FP were supported by the Novo Nordisk Foundation grant NNF20SA0035590 and NNF0069780. AH acknowledges funding by EMBO (ALTF 179-2021) and the European Crohńs and Colitis Organization (PROP-1495). The Novo Nordisk Foundation Center for Stem Cell Medicine was supported by a Novo Nordisk Foundation grant (NNF21CC0073729). MM is funded by the Max Planck Society for the Advancement of Science.

## Conflicts of interest

CS lectures for MSD and Janssen-Cilag and received a research grant from Takeda. MM is an indirect shareholder in Evosep.

---

**Figure S1.**
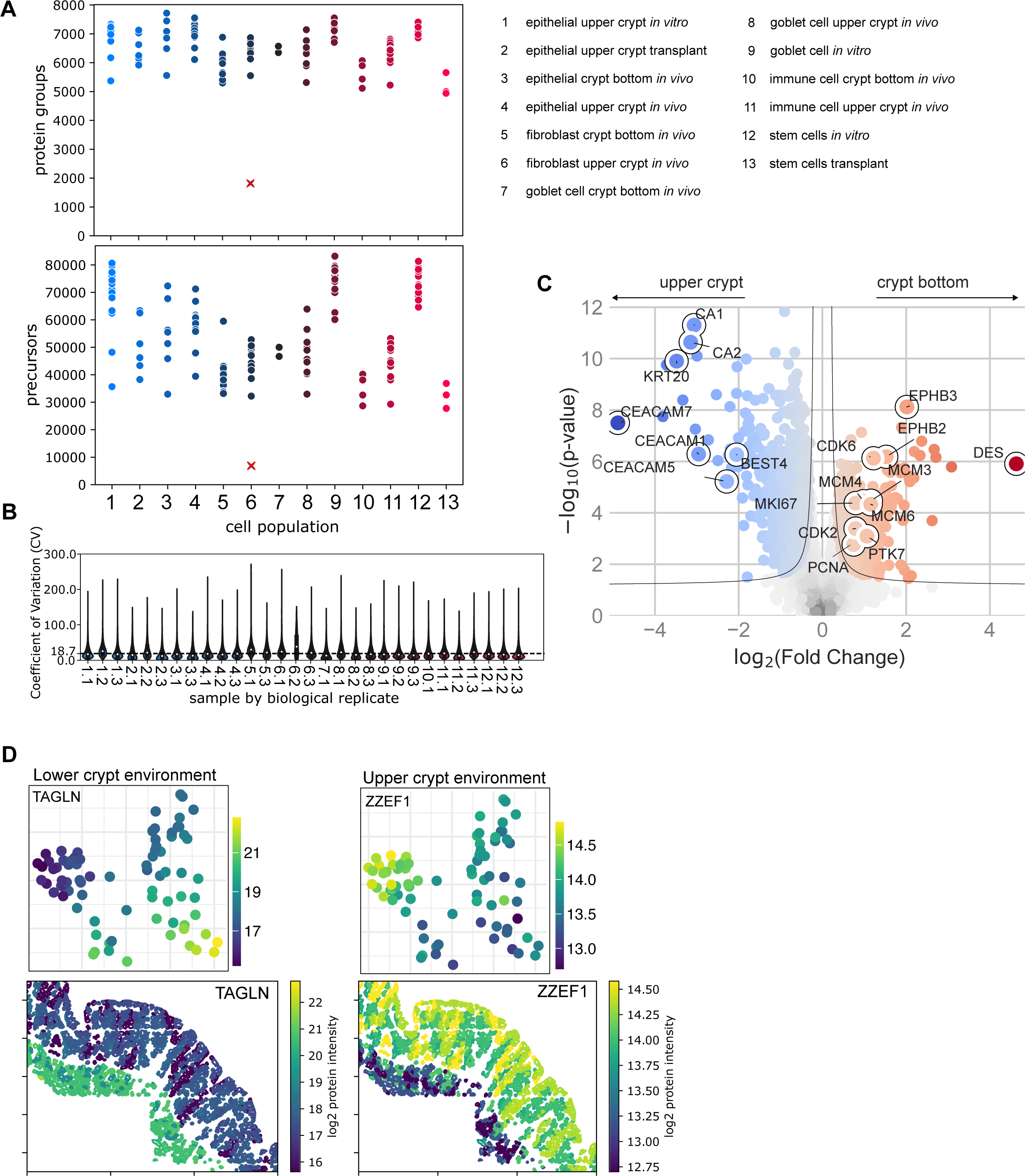
A Number of identified proteins and precursors per sample. **B** Coefficient of variation of technical replicates. **C** Volcano plot comparing epithelial cells from the crypt bottom and upper crypt *in vivo*. **D** Protein abundance and spatial distribution of TAGLN and ZZEF1, which are differentially abundant in the crypt bottom versus upper crypt region in the colon mucosa across different cell types.

**Figure S2.**
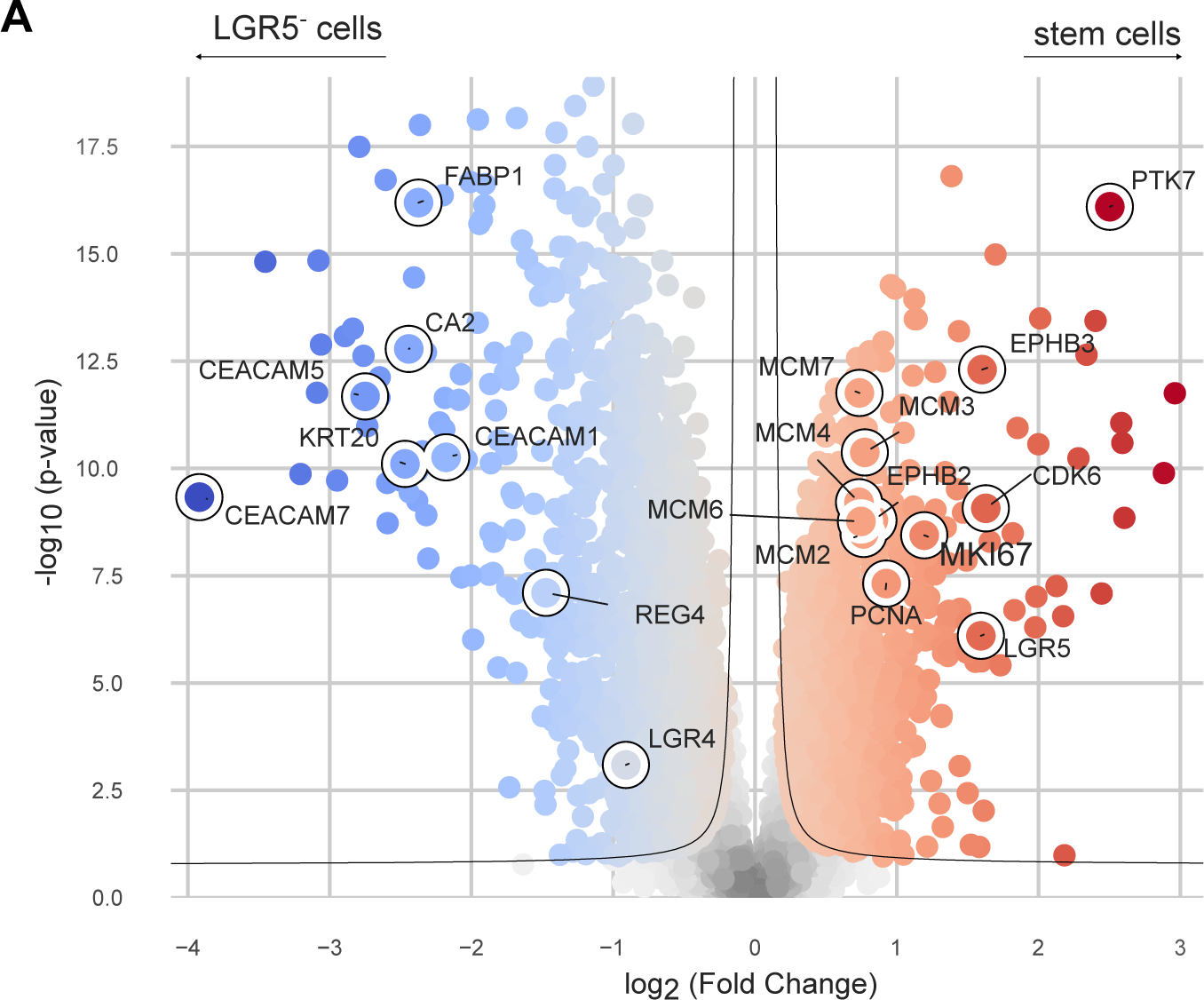
A Volcano plot of stem cells (LGR5-TdTomato^+^) and LGR5-TdTomato^-^ cells in organoids *in vitro*.

**Figure S3.**
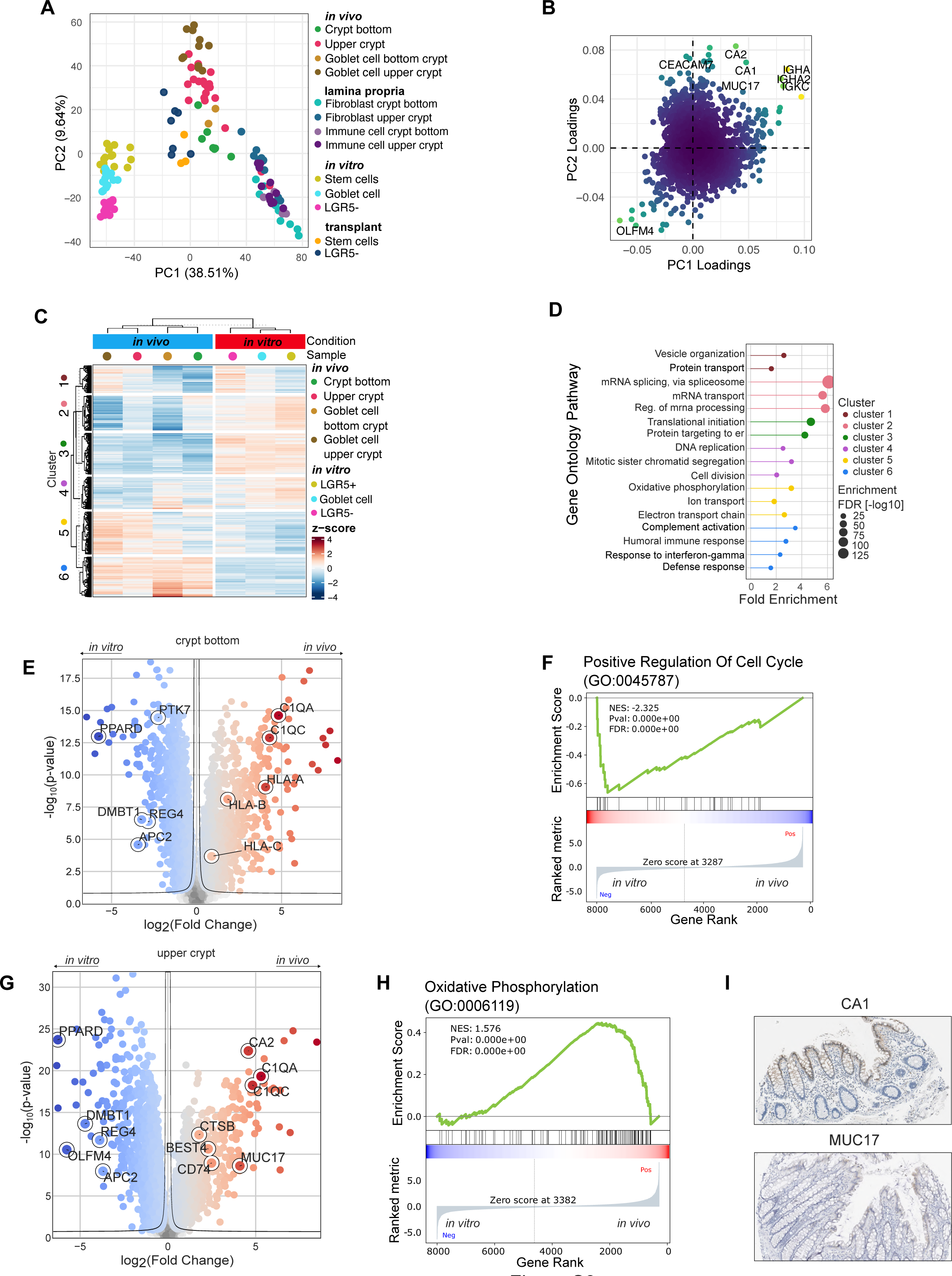
A PCA of the top 3000 most varying proteins across samples in *in vitro*, transplant, and *in vivo*. **B** Loadings describing proteins driving the PCA in S3A. CA1, CA2, MUC17 and CEACAM7 are strongly associated with a crypt top *in vivo* colonocyte phenotype. **C** Heatmap of significantly changed proteins between epithelial cells *in vitro* and *in vivo*. **D** Pathway enrichments of proteins in clusters of Fig S3C. **E** Volcano plot of stem cells *in vitro* vs crypt bottom epithelial cells *in vivo*. **F** Gene Set Enrichment Analysis (GSEA) of the Gene Ontology term “positive regulation of cell cycle” on protein differences of S3E. **G** Volcano plot of epithelial cells in the upper crypt *in vitro* vs *in vivo*. **H** GSEA of the Gene Ontology term “oxidative phosphorylation” on protein differences of S3G. **I** Staining for CA1 and MUC17 in the human colon mucosa from the Human Protein Atlas^43^.

**Figure S4.**
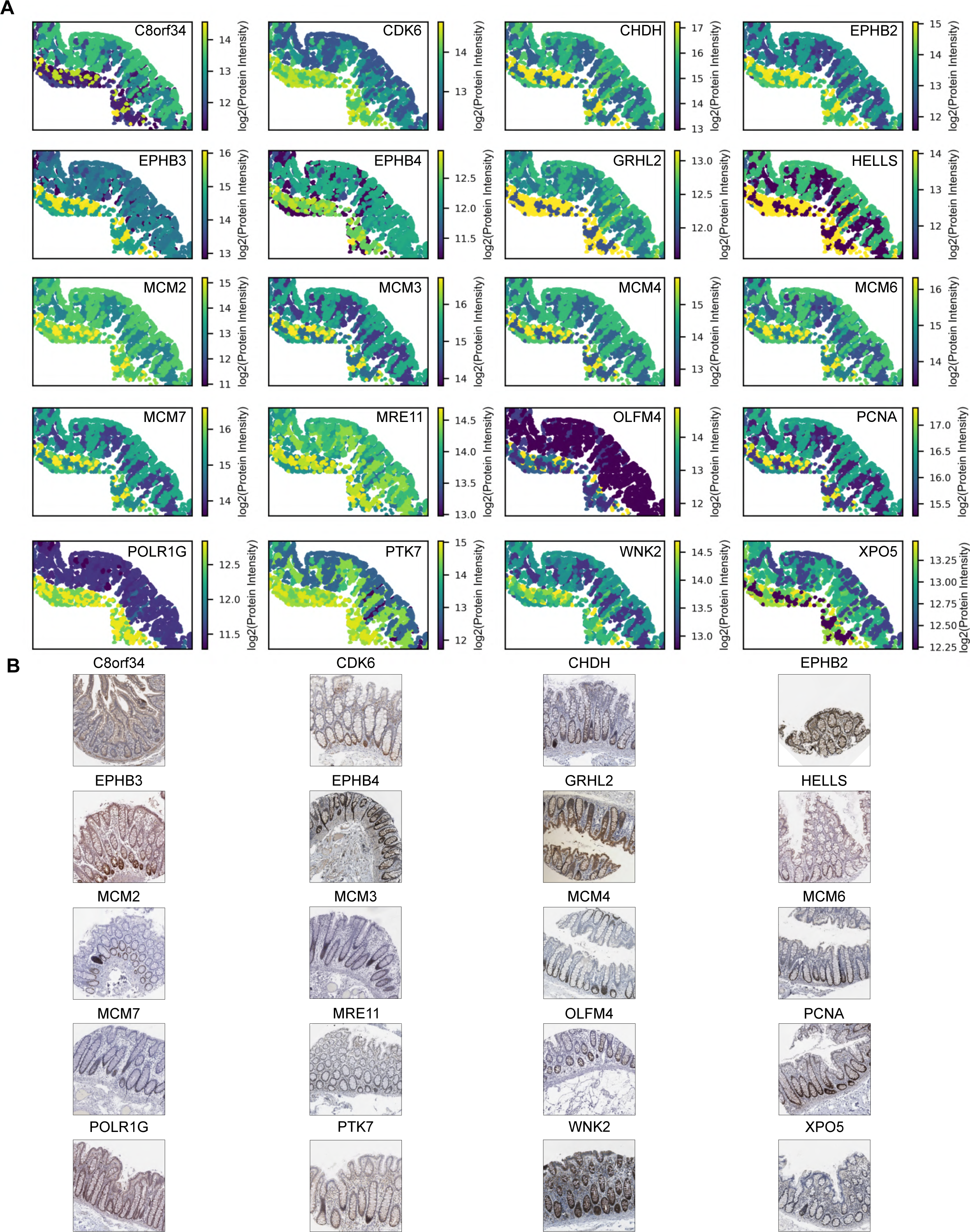
A Spatial distribution of proteins that were identified in the colon stem cell signature. **B** Immunohistochemistry staining from the Human Protein Atlas^43^ in human colon of proteins that were identified as potential colon stem cell markers.

**Figure S5.**
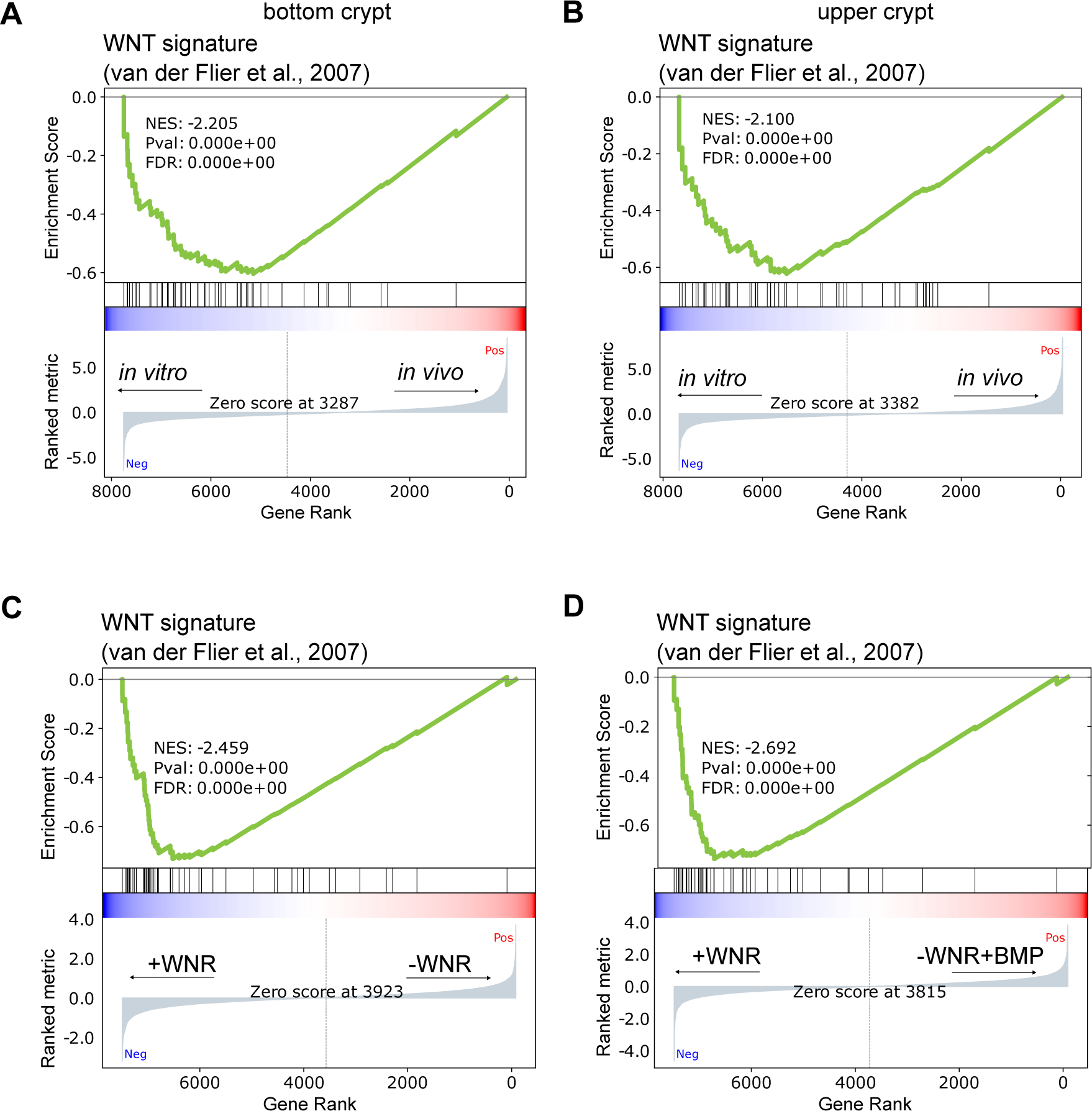
A Gene set enrichment analysis of WNT signature proteins^51^ in crypt bottom IECs *in vivo* versus stem cells *in vitro*, **B** in upper crypt IECs *in vivo* versus LGR5-TdTomato^-^ cells *in vitro*, **C** organoids grown under -WNR versus +WNR conditions and **D** organoids grown under -WNR+BMP and +WNR conditions.

